# Early modern human dispersal from Africa: genomic evidence for multiple waves of migration

**DOI:** 10.1101/022889

**Authors:** Francesca Tassi, Silvia Ghirotto, Massimo Mezzavilla, Sibelle Torres Vilaça, Lisa De Santi, Guido Barbujani

**Author notes:** Correspondence and requests for materials should be addressed to Guido Barbujani. These authors contributed equally to this work (Francesca Tassi, Silvia Ghirotto). Francesca Tassi; Silvia Ghirotto; Massimo Mezzavilla; Sibelle Torres Vilaça; Lisa De Santi; Guido Barbujani.

## Abstract

**Background:** Anthropological and genetic data agree in indicating the African continent as the main place of origin for modern human. However, it is unclear whether early modern humans left Africa through a single, major process, dispersing simultaneously over Asia and Europe, or in two main waves, first through the Arab peninsula into Southern Asia and Oceania, and later through a Northern route crossing the Levant.

**Results:** Here we show that accurate genomic estimates of the divergence times between European and African populations are more recent than those between Australo-Melanesia and Africa, and incompatible with the effects of a single dispersal. This difference cannot possibly be accounted for by the effects of hybridization with archaic human forms in Australo-Melanesia. Furthermore, in several populations of Asia we found evidence for relatively recent genetic admixture events, which could have obscured the signatures of the earliest processes.

**Conclusions:** We conclude that the hypothesis of a single major human dispersal from Africa appears hardly compatible with the observed historical and geographical patterns of genome diversity, and that Australo-Melanesian populations seem still to retain a genomic signature of a more ancient divergence from Africa

## Background

Anatomically modern humans (AMH), defined by a lightly built skeleton, a large brain, reduced face and prominent chin, first appear in the East African fossil record around 200,000 years ago [1, 2]. There is a general consensus that, while dispersing from there, they largely replaced preexisting archaic human forms [3]. Recent DNA studies also suggest that the replacement was not complete, and there was a limited, but nonzero, interbreeding with Neandertals [4], Denisovans [5], and perhaps other African forms still unidentified at the fossil level [6, 7]. As a result, modern populations might differ for the amount of archaic genes incorporated in their gene pool, which are eventually expressed and may result in phenotypic differences affecting, for example, the immune response [8], or lipid catabolism [9].

Although the general picture is getting clearer, many aspects of these processes are still poorly understood, starting from the timing and the modes of AMH dispersal. The main exit from Africa, through the Levant, has been dated around 56,000 years ago [10, 11]. However, morphologic [12, 13], archaeological [14] and genetic [13, 15-20] evidence suggest that part of the AMH population might have dispersed before that date, possibly by a Southern route into Southern Asia through the horn of Africa and the Arab peninsula.

Regardless of whether there was a single major expansion or two, several DNA studies clearly showed that genetic diversity tends to decrease [21, 22] and linkage disequilibrium to increase [23, 24] at increasing distances from Africa. This probably means that, as they came to occupy their current range, AMH went through a series of founder effects [25, 26]. These results offer an excellent set of predictions which we used in the present study to test whether current genomic diversity is better accounted for by processes involving a Single major Dispersal (hereafter: SD) or Multiple major Dispersals (hereafter: MD) from Africa.

One preliminary problem, however, is how to select the appropriate populations for informative comparisons. The details of the dispersal routes, and the relationships between fossils and contemporary populations, are all but established. Whereas Europeans are consistently regarded as largely derived from the most recent African exit in all relevant studies, opinions differ as for many aspects of the peopling of Asia [12-19], with many populations also experiencing complex demographic histories involving admixture, as suggested by both ancient [27] and modern [28-31] DNA evidence. To obtain insight into the past history of Eurasian populations we analyzed genome-wide autosomal single nucleotide polymorphisms (SNPs) from 71 worldwide populations (Additional File 1 and Additional File 2). In what follows, a number of preliminary analyses allowed us to quantify the extent and the pattern of admixture and gene flow in our data, thus making it possible to identify a subset of Far eastern populations which, under the MD model, may safely be regarded as deriving from the oldest expansion.

This way, we could address two questions, related, respectively, with the historical and geographical context of the dispersal process, namely: (1) are separation times between non-African and African populations the same (as expected under SD), or is there evidence of a longer separation between Far Eastern and Africans than between Europeans and Africans (as expected under MD)? And (2) which geographical migration routes were followed by first humans outside Africa?

## Materials and Methods

### Populations and markers

We combined genomic data from several published datasets: the Human Genome Diversity Cell Line Panel [32] (n = 40 samples from 10 populations genotyped on Affymetrix GeneChip Human Mapping 500 K Array Set), Pugach et al (2013) [33] (n= 117 samples from 12 populations genotyped on an Affymetrix 6.0 array), Reich et al (2009) [34] (n = 56 samples from 11 populations genotyped on an Affymetrix 6.0 array), Reich et al (2011) [5] (n = 509 samples from 13 populations genotyped on an Affymetrix 6.0 array), Xing et al (2009) [35] (n = 243 samples from 17 populations genotyped on one array (version NspI) from the Affymetrix GeneChip Human Mapping 500K Array set), Xing et al (2010)[36] (n = 165 samples from 8 populations genotyped on an Affymetrix 6.0 array), (Additional File 1 and Additional File 2).

We devised a careful strategy to combine the seven datasets genotyped with different platforms according to different protocols developing a pipeline built on Perl. First, for each dataset, we checked for the presence of old rs ids, if necessary changing them with the new ones. Then, we looked for the SNPs shared among all datasets and we mapped the genome positions of these variants to the human reference genome, build hg18 (NCBI 36).

When merging data from different SNP-chip versions, strand identification can be ambiguous, possibly leading to mistakes in identifying the right alleles for A/T and G/C SNPs (as also reported in the PLINK tool documentation [37]). Thus, to preserve as much genetic information as possible, we selected from each dataset only these ambiguous SNPs and we used the information contained in Affymetrix Annotation file to evaluate the strand polarity used to define each allele. We considered each dataset separately and we annotated the SNPs on the plus strand, flipping only the proper SNPs. We checked the reliability of this conversion process comparing the allele frequencies for these SNPs in specific populations typed in more than one dataset (i.e Besemah, CEU, Onge), so as to verify the consistency of the frequency spectrums between the different datasets. Once these ambiguities have been resolved, with the PLINK v 1.07 software [37] we merged progressively the datasets selecting, from each one, just the individuals from populations of our interest and flipping SNPs discordant for strand.

Using the same software, we selected only the autosomal SNPs with genotyping success rate >98% and minor allele frequency (MAF) >0.01. We identified cryptic relatedness amongst samples computing identity-by-descent (IBD) statistic for all pairs of individuals, as unmodeled excess of genetic sharing would violate sample independence assumption of downstream analyses. When pairs of individuals showed a Pi-Hat value > 0.3, we removed the individual with the lowest genotyping rate. We did not apply this screening procedure for the South-East Asia and Oceania samples, since they come from populations with extremely low effective sizes, where a certain degree of random inbreeding is inevitable [38]. To determine whether there were genetic outliers within each population, we conducted in PLINK a “distance to the nearest neighbor analysis” (--neighbor option). Within each population, the measure of similarity in terms of identity by state (IBS) between each individual and their nearest neighbor was calculated and transformed into a Z-score. Z score distributions were examined from the first to the fifth neighbor. Outliers were identified by an extremely negative Z-score produced by *low* allele sharing with their nearest neighbor and were then dropped from the population. We grouped populations according to ethnological and linguistic information; the final dataset is shown in Figure 1a.

**Figure 1.**
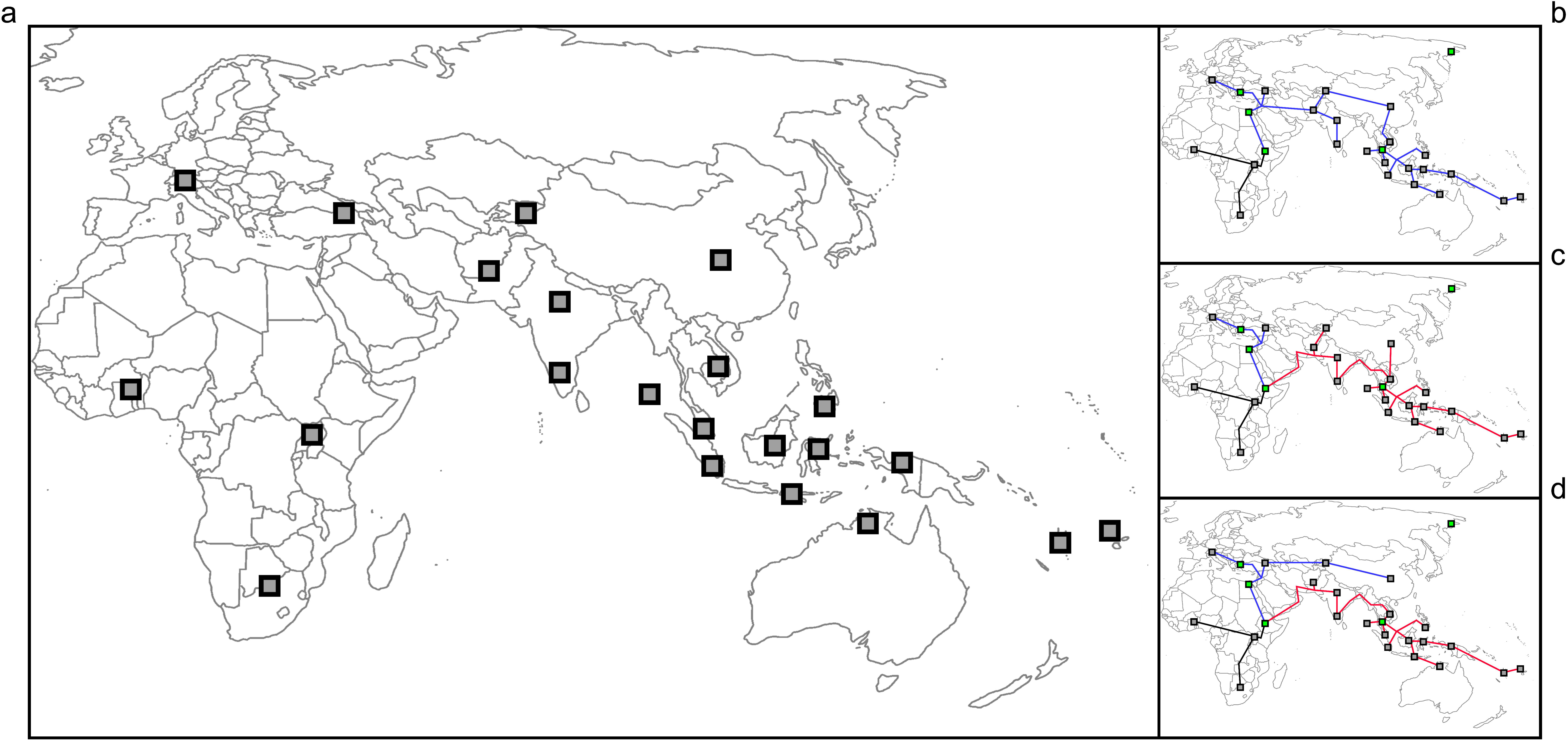
Geographic location of the 24 metapopulations analyzed (a) and geographical models of African dispersal (b, c, d). Metapopulations, each derived from the merging of genomic data from several geographically or linguistically-related populations, are South, East and West Africa, Europe, the Caucasus, South, East, West and Central Asia, North and South India, plus three Negrito (Onge, Jehai and Mamanwa) and ten Oceanian populations; the final dataset comprised 1,130 individuals. Under model 1, a SD model (b), all non-African populations are descended from ancestors who left Africa through the same, Northern route[3]. Model 2 (c) and Model 3 (d) are MD models assuming, prior to dispersal across Palestine, another exit through the Arab Peninsula and the Indian Subcontinent; under Model 2 all Asian and Western Oceanian populations derive from this earlier expansion[12], whereas under Model 3 only the populations of Southeast Asia and Western Oceania derive from the earlier expansion [16].

To visualize the genetic relationships between such populations, we performed a Principal Component Analysis using the R [39] SNPRelate package.

### Population structure analysis

Individual genotypes were clustered, and admixture proportions were inferred, by the algorithm embedded in the software ADMIXTURE, based on the principle of maximum likelihood[40]. This method considers each genotype as drawn from an admixed population with contributions from *k* hypothetical ancestral populations. Because this model assumes linkage equilibrium among markers, we checked with the PLINK v1.07 tool [37] that the set of SNPs we used did not show a level of Linkage Disequilibrium higher than r^2^=0.3; this way, in the pruned dataset 54,978 markers were retained. The optimal value of *k* was evaluated through a cross-validation procedure, testing values from *k*=2 to *k*=14, thus identifying the number of ancestral populations for which the model had the best predictive accuracy. We then ran an unsupervised analysis, assuming a number of ancestral admixing populations from *k*=2 to *k*=7. The proportion of the individuals’ genome belonging to each ancestral population was calculated for each k value from 5 independent runs, then combined by the software CLUMPP [41] and plotted by the software *Distruct* [42].

### Discriminant Analysis of Principal Components

In addition to ADMIXTURE, to identify and describe clusters of genetically related individuals we used a Discriminant Analysis of Principal Components (DAPC) [43] implemented in the R [39] package adegenet ver. 1.3-9.2 [44].DAPC methods allow one to assess the relationships between populations overlooking the within-group variation and summarizing the degree of between group variation. Being a multivariate method, DAPC is suitable for analysing large numbers of genome-wide SNPs, providing assignment of individuals to different groups and an intuitive visual description of between-population differentiation. Because it does not rely on any particular population genetics model, DAPC is free of assumptions about Hardy-Weinberg equilibrium or linkage equilibrium [43], and so we could use the full set of 96,156 SNPs for this analysis.

By the function *find.clusters*, we determined the most likely number of genetic clusters in our dataset, using all principal components (PCs) calculated on the data. The method uses a K-means clustering of principal components [10] and a Bayesian Information Criterion (BIC) approach to assess the best supported number of clusters.

Then, we determined the optimal number of principal components (PCs) to retain to perform a discriminant analysis avoiding unstable (and improper) assignment of individuals to clusters. It is worth noting that, unlike K-means, DAPC can benefit from not using too many PCs: retaining too many components with respect to the number of individuals can lead to over-fitting and instability in the membership probabilities returned by the method.

### Population divergence dates

The divergence times between populations (*T*), was estimated from the population differentiation index (*F*_*ST*_) and the effective population size (*Ne*). F_ST_ is the proportion of the total variance in allele frequencies that is found between groups and it was calculated between pairs of populations for each SNP individually under the random population model for diploid loci, as described by Weir and Cockerham [45], and then averaged over all loci to obtain a single value representing pairwise variation between populations. Under neutrality, the differences between populations accumulate because of genetic drift, and so their extent depends on two quantities: it is inversely proportional to the effective population sizes (*N*_*e*_) and directly proportional to the time passed since separation of the two populations (*T*).

Therefore, to estimate *T* from genetic difference between populations, independent estimate of *N*_*e*_ are needed; for this purpose we focused on the relationship between *N*_*e*_ and the level of linkage disequilibrium within populations. Indeed, levels of *LD* also depend on *Ne*, and on the recombination rate between the SNPs considered [46], with *LD* between SNPs separated by large distances along the chromosome reflecting the effects of relatively recent *N*_e_, whereas *LD* over short recombination distances depending on relatively ancient *N*_e_ [47]. Thus, we estimated *LD* independently in each population using all polymorphic markers available for that population (MAF > 0.05), from a minimum of ∼ 90,000 SNPs in Polynesia to a maximum of ∼ 370,000 SNPs in North India. This way, we also reduced the impact of ascertainment bias, i.e. the bias due to the fact that most SNPs in the genotyping platforms were discovered in a single (typically European) population [48].

We assigned to each SNP a genetic map position based on HapMap2 (Release #22) recombination data, and for each pair of SNPs separated by less than 0.25 cM we quantified *LD* as r^2^_LD_ [49] or as σ^2^_LD_ [50] (hereafter: *ρ*). All the observed *ρ* values were then binned into one of 250 overlapping recombination distance classes. Pairs of SNPs separated by less than 0.005 cM were not considered in the analysis, since at these very short distances gene conversion may mimic the effects of recombination [46]. We also adjusted the *ρ* value for the sample size using 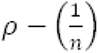 [46]. Finally, we calculated the effective population size for each population in each recombination distance class as

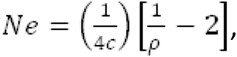

corresponding to the effective population size 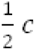 generations ago, where *c* is the recombination distance between loci, in Morgans [47, 51, 52]. Finally, the long-term *N*_e_ for each population was calculated as the harmonic mean of *N*_e_ over all recombination distance classes up to 0.25 cM. The confidence intervals of these *N*_e_ values were inferred from the observed variation in the estimates across chromosomes.

Based on the independently-estimated values of *N*_e_, we could then estimate *T* as

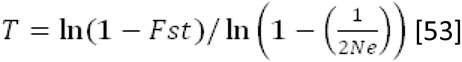

where time is expressed in generations.

All procedures were performed by in-house developed software packages, NeON [54] and 4P [55].

### Simulations

To understand whether the divergence times estimated were compatible with a SD model, we used a neutral coalescent approach to simulate genetic polymorphism data under the infinite-sites model of mutation. We simulated data representing 1Mb chromosome segments in two populations according to the demographic scenario shown in Additional File 3 using the coalescent simulator *ms* [56]. We assumed an ancestral population with an initial Ne= 10,000. At t = T, the population splits into two populations. Population_2a’s Ne remains constant, population_2b has a 50% reduction in Ne followed by an exponential growth, representing the genetic bottleneck experienced by populations dispersing out of Africa. In all simulations the scaled mutation rate (θ), and the scaled population recombination rate (ρ) were fixed at 400. For a sequence length of 1Mb and an effective population size of Ne= 10,000 these parameters correspond to a mutation rate of 10^−8^ and a recombination rate of 1 cM/Mb.

To account for the uncertainty in the estimates of both the timing of the process and the effective population sizes, according to Ref. [57-59], this model was simulated considering 4 different separation times (T) (between 40,000 and 70,000 years ago, in steps of 10,000 years) and 6 estimates of the actual effective size for population_2b (between 3,000 and 8,000, in steps of 1,000). For each of the 24 simulation conditions, 1,000 independent datasets were simulated and then analyzed according to the following procedure:

1. A sample of 50 individuals (i.e. 100 chromosomes) was randomly selected from each population. The simulated genetic data were single nucleotide polymorphisms (SNPs) segregating within the two populations.
2. We converted the *ms* [56] output file to PLINK format [37].
3. Any SNPs with a minor allele frequency (MAF) less than 0.05 were removed from the datasets.
4. We estimated the population differentiation index, effective population size and divergence time between the two simulated populations following the same procedures used for the observed data and detailed above.
5. Estimators were calculated for each 1,000 independent replications.

### Possible effects of a Denisovan admixture in Melanesia

To rule out the possibility that the divergence time estimated between Africans and New Guinea/Australia samples could reflect, largely or in part, admixture between the Denisovan archaic human population from Siberia[60] and the direct ancestor of Melanesians, we removed from our dataset the variants could be regarded as resulting from such a process of introgression. These SNPs would carry the derived state in the archaic population and in the New Guinean/Australian samples, while being ancestral in East Africans and Europeans (i.e. those populations that did not show any signal of introgression from Denisova [5, 60]).

Using the *VCF tools*[61] we extracted our 96,156 SNP from the high coverage Denisovan genome. We then removed from these data all transitions SNPs (C/T and G/A) because in ancient DNA these sites are known to be prone to a much higher error rate than the transversions [5]. Then, we selected the sites meeting the following set of criteria:

- the site has human-chimpanzee ancestry information;
- himpanzee ancestral allethe human-chimpanzee ancestral allele matches one of the two alleles at heterozygous sites;
- Denisova has at least one derived allele;
- New Guineans and Australians have at least one derived allele;
- in East African and European individuals the ancestral allele is fixed;

When the ancestry information was missing (1,438 SNPs), to define the ancestral state, we used the East African individuals selecting the SNPs where East Africans were homozygous and considering those as ancestral.

Once we had thus identified a subset of sites putatively introgressed from Denisova, we removed them from the dataset. The remaining 80,621 SNPs were used to compute the pairwise F_ST_[45] values and to infer the divergence time between populations, as described above.

### Testing for the effect of recent gene flow

Using *TreeMix*, we inferred from genomic data a tree in which populations may exchange migrants after they have split from their common ancestor, thus violating the assumptions upon which simple bifurcating trees are built [62]. This method first infers a maximum-likelihood tree from genome-wide allele frequencies, and then identifies populations showing a poor fit to this tree model; migration events involving these populations are finally added. This way, each population may have multiple origins, and the contributions of each parental population provide an estimate of the fraction of alleles in the descendant population that originated in each parental population.

Allele frequencies for the *TreeMix* analysis were calculated by PLINK tool [37], after pruning for linkage disequilibrium as we did for ADMIXTURE analysis. We modelled several scenarios allowing a number of migration events from 0 to 6, and stopping adding a migration when the following event did not increase significantly the variance explained by the model. The trees were forced to have a root in East Asia, and we used the window size of 500 (-k option).

### Geographical patterns of dispersal

We developed explicit geographic models of demographic expansion, and looked for the model giving the closest association between genomic and geographical distances. In all cases, migration routes were constrained by 5 obligatory waypoints, identified in Ref. [26] and accepted by several successive studies (see e.g Ref.[13]). In addition, because of some inconsistencies in the definition of the geographic regions affected by the two waves of migration under MD [12, 14, 16], and of the ambiguity introduced by the previously described episodes of admixture, two different models of MD were considered. Under Model 1, a SD model, anatomically modern humans left Africa through Palestine and dispersed into both Europe and Asia (Figure 1b). Model 2 assumes, prior to the dispersal across Palestine, another exit through the Arab Peninsula and the Indian Subcontinent, all the way to Melanesia and Australia; according to this model, based on skull morphology [12] all Asians populations are derived from this earlier expansion (Figure 1c). On the contrary, under Model 3 only the populations of Southeast Asia and Oceania are derived from the earlier expansion, whereas Central Asian populations are attributed to the later African dispersal [16] (Figure 1d).

To obtain a realistic representation of migrational distances between populations, we did not simply estimate the shortest (great-circle) distances between sampling localities. Rather, we modelled resistance to gene flow, based on the landscape features known to influence human dispersal. We used a resistance method from the circuit theory implemented in the software Circuitscape v.3.5.2 [63], starting from the landscape information in Ref [64] and referring to the distribution of land masses at the last glacial maximum. Next, we added data about altitude and river presence from the Natural Earth database. Each area of the map was assigned a resistance value (*rv*) by the Reclassify tool in ArcGIS 10 (ESRI; Redlands, CA, USA), as follows: mountains higher than 2,000 m: *rv*=100; land or mountains below 2,000 m: *rv*=10; rivers: *rv*=5, oceans: NoData (absolute dispersal barrier); narrow arms of sea across which prehistoric migration is documented: *rv*=10. The low *rv* for rivers reflects the human tendency to follow, whenever possible, water bodies in their dispersal (see e.g. Ref. [65]).

Under the SD model we hampered movement from Arabia to India (*rv*=100), hence preventing the dispersal along the Southern route; under the MD models, we created a buffer of low resistance value (*rv*=1) along the Southern route. For all models we then estimated least-resistance distances between the populations analyzed, when applicable going through Addis Ababa, chosen as a starting point for the African expansion [26]. The final output was then exported in Google Earth where geographic distances were expressed in kilometers.

We evaluated by partial Mantel tests [66] the correlation between genomic (*F*_ST_) and geographic distances, while holding divergence times constant. This way we could control for the drift effects, due to the fact that populations separated at distinct points in time and space.

## Results

### Genomic structure of Old World populations

We assembled genome-wide SNP data from the literature obtaining information on 71 population samples sharing, after cleaning and integration, 96,156 autosomal SNPs. By merging samples from adjacent geographical regions and with similar linguistic affiliations, we organized the data in 24 meta-populations; the final dataset comprised 1,130 individuals (Figure 1a and Additional File 2).

As a preliminary step, we visualized by Principal Component Analysis the genetic relationships between such populations, as inferred from these autosomal SNPs (Fig. 2). The first two principal components, accounting respectively for 8.4% and 4.3% of the total genetic variance, show that the populations we grouped in meta-populations do cluster together genetically. In addition, genetic relationships largely correspond to geographical distances, with Eurasian populations separated from the African ones along the axis represented by PC1, and forming an orderly longitudinal cline, all the way from Europe to East Asia and Oceania, along the PC2 axis.

**Figure 2.**
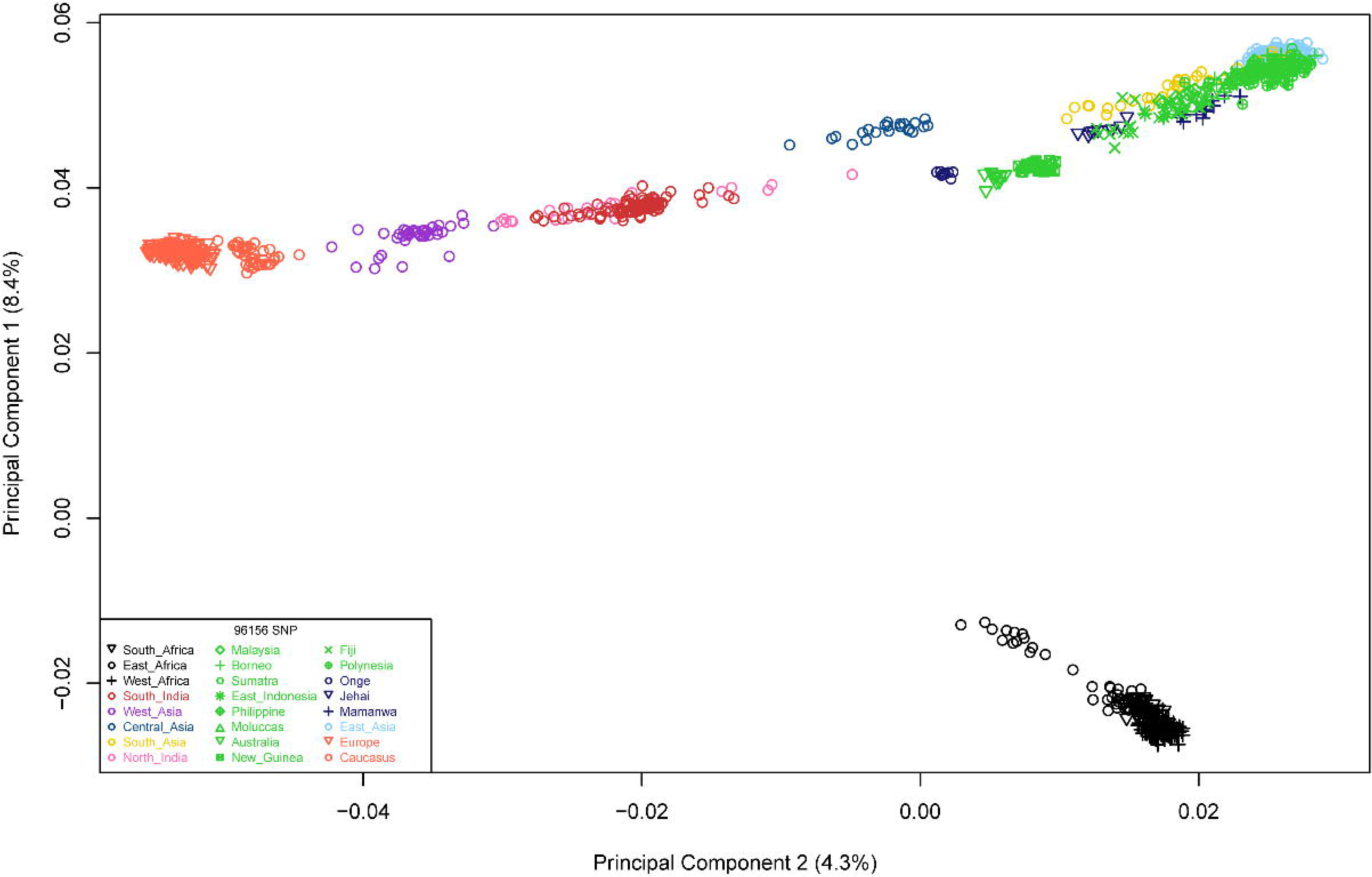
Results of the Principal Component Analysis. Each symbol corresponds to an individual genotype; the first two principal components account for 12.7% of the global variation in the data. Here and in all figures, different colours represent different geographical regions.

Then, to further investigate the worldwide genomic structure, we applied the unsupervised ancestry-inference algorithm of the ADMIXTURE software [40]. After identifying k=6 as the most supported number of ancestral populations (Additional File 4), we explored the results for *k* = 2-7 ancestral populations. As the number of ancestral clusters increased, we observed the emergence of several well-supported population-specific ancestry clusters (Figure 3). At *k* = 2, the ancestry assignment differentiated between African (blue) and non-African (yellow) populations; *k*=3 further distinguishes Europeans from Asians (orange); *k*=4 identifies an Australo-Melanesian component (green) within the Asian cluster; at *k*=5 the additional component is mainly associated with the Indian subcontinent (red); the same is the case at *k*=6 for Polynesia and Fiji (pink) and at *k*=7 for many island communities of Southeast Asia and Oceania (purple). Remarkably, some populations show more than 99% contribution from the same ancestral population along different *k* values (e.g. West Africa, Europe, New Guinea), whereas other populations include several individuals with an apparently admixed genomic background, possibly resulting from successive gene flow (e.g. back migration from Europe to Northeast Africa [67]). A Discriminant Analysis of Principal Components (DAPC)[43] led to essentially the same conclusions as ADMIXTURE. We found *k*=6 to be the best supported model (Additional File 5) and therefore used this value in the DAPC. Additional File 6a shows that the main populations are distinguishable, and most individuals from the same population tend to fall in the same cluster. In the scatter plot the first two axes revealed three major clusters within the six supported by the *k*= 6 model (Additional File 6b). They included (i) the three African population, (ii) most populations from Asia, and (iii) populations from Europe and Caucasus and from India and West-Asia. This clustering pattern is also observed in Admixture analysis with *k*= 6 (Figure 3). Interestingly, in the Asian group the DAPC is able to distinguish three different clusters: one represented by individuals from Australia and New Guinea (in green color), one by the populations showing at least 30% of the green Admixture component at *k*=5 (in pink color), and one by other populations from Asia.

**Figure 3.**
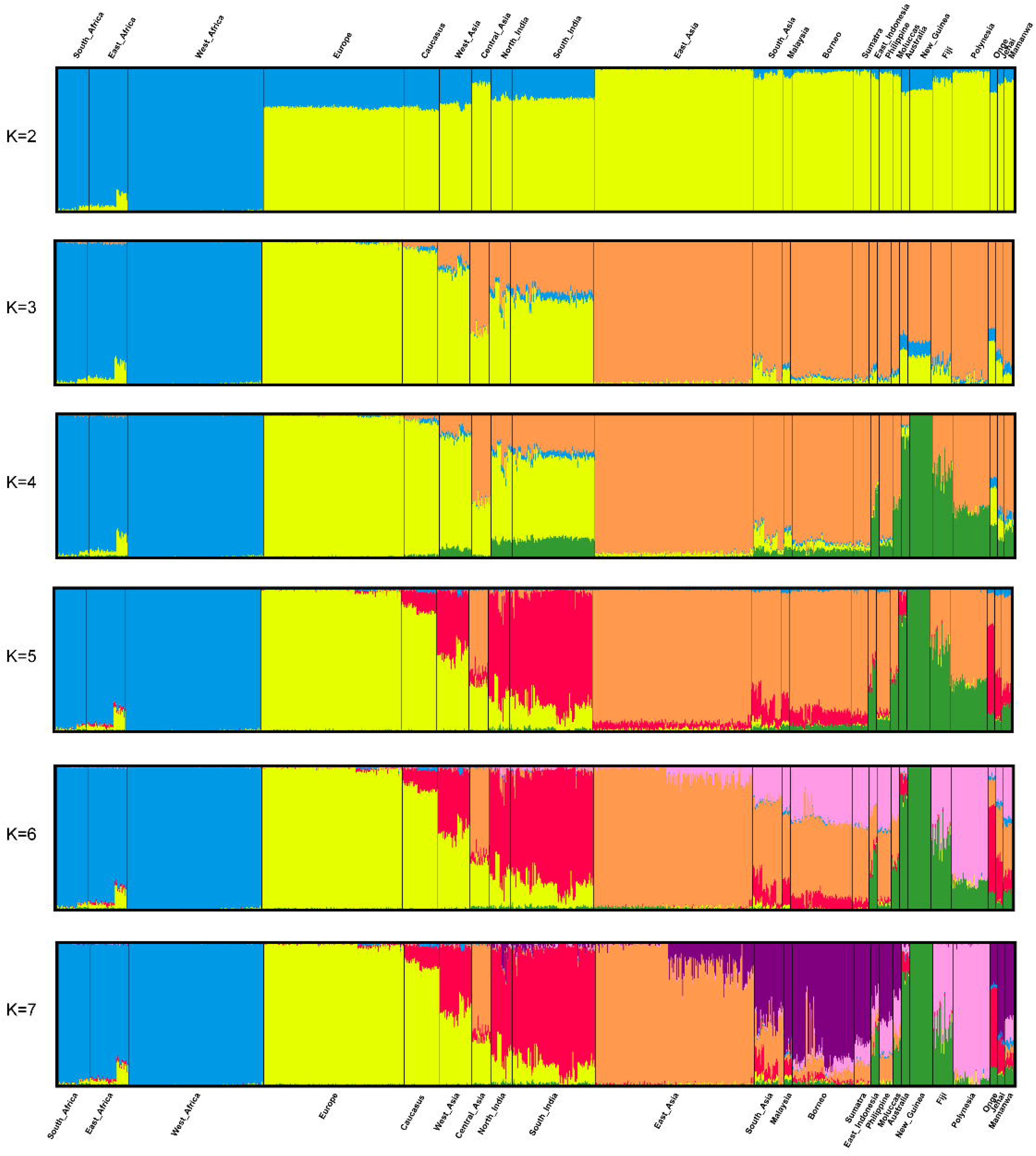
Admixture analysis of 1130 individuals in 24 populations from Africa, Eurasia and Western Oceania. Each individual genotype is represented by a vertical column, the colors of which correspond to the inferred genetic contributions from *k* putative ancestral populations. The analysis was run for 2 ≤ *k* ≤ 7

### Population divergence dates

There is a clear geographical structure in the data, which in principle allows one to test for the relative goodness of fit of the two models. The SD model implies that the separation time from Africa of all samples should be the same, whereas significantly larger times of separation are expected under the MD model for the Eastern most than for the European populations. To answer, we calculated the divergence time between each out of Africa population and East Africa, South Africa and West Africa populations as described in the Materials and Methods section, obtaining an independent estimate of Ne from LD which allowed us to tell apart the effects of population sizes and separation times in the observed F_ST_ values.

The three African populations show the largest long-term population sizes (calculated as the harmonic mean of Ne values) and a constant declining trend through time, whereas Eurasian populations (and more markedly the Asian ones) tend to increase in size, especially in the last 10,000 years. Australians and New Guineans (represented in green in the Admixture analysis at *k* ≥ 4), generally maintain a constant size until present times, whereas the Negrito populations show low and declining sizes. In general, these results were not surprising, but the fact we obtained them suggests that the procedure followed is by and large accurate, and therefore that the estimated average *N*_e_s are plausible. The values obtained using the two estimators of LD (*r*^2^ and σ^2^, see Materials and Methods section) gave similar results (Additional File 7 and Additional File 8).

From the pairwise *F*_ST_ values estimated over all loci (Additional File 9), and now considering the independently-estimated values of *N*_e_, we could infer the divergence times between populations (Table 1 and Additional File 10). The average separation times from the East African populations, i.e. those located in the most plausible site of departure of AMH expansions [26] (Table 1) are distributed along a range spanning from 60K to 100K years ago. Extreme divergence values were observed for Europe and the Caucasus on the one hand, and for Australia and New Guinea on the other, respectively at the lower and the upper tails of the distribution. Even considering the full range of uncertainty around these estimates (95% of the confidence interval) we observed no overlap, with Europe having an upper confidence limit 77/71K years ago (depending on the LD measure used, respectively the *r*^2^ and σ^2^ statistic) and Australia having a lower confidence limit 88/80K years ago. Because we kept into consideration the effects of *N*_e_ in the estimation procedure, this difference cannot possibly be accounted for by the different impact of genetic drift upon these populations, and supports a rather complex “Out of Africa” scenario, suggesting at least two main phenomena of AMH dispersal from Africa. The Australo-Melanesian populations, i.e. Australians and New Guineans, with their early separation times from East Africa, may be regarded as the putative descendants of an early dispersal process, whereas the status of most Asian populations would seem, at this stage of the analysis, unclear.

**Table 1.**
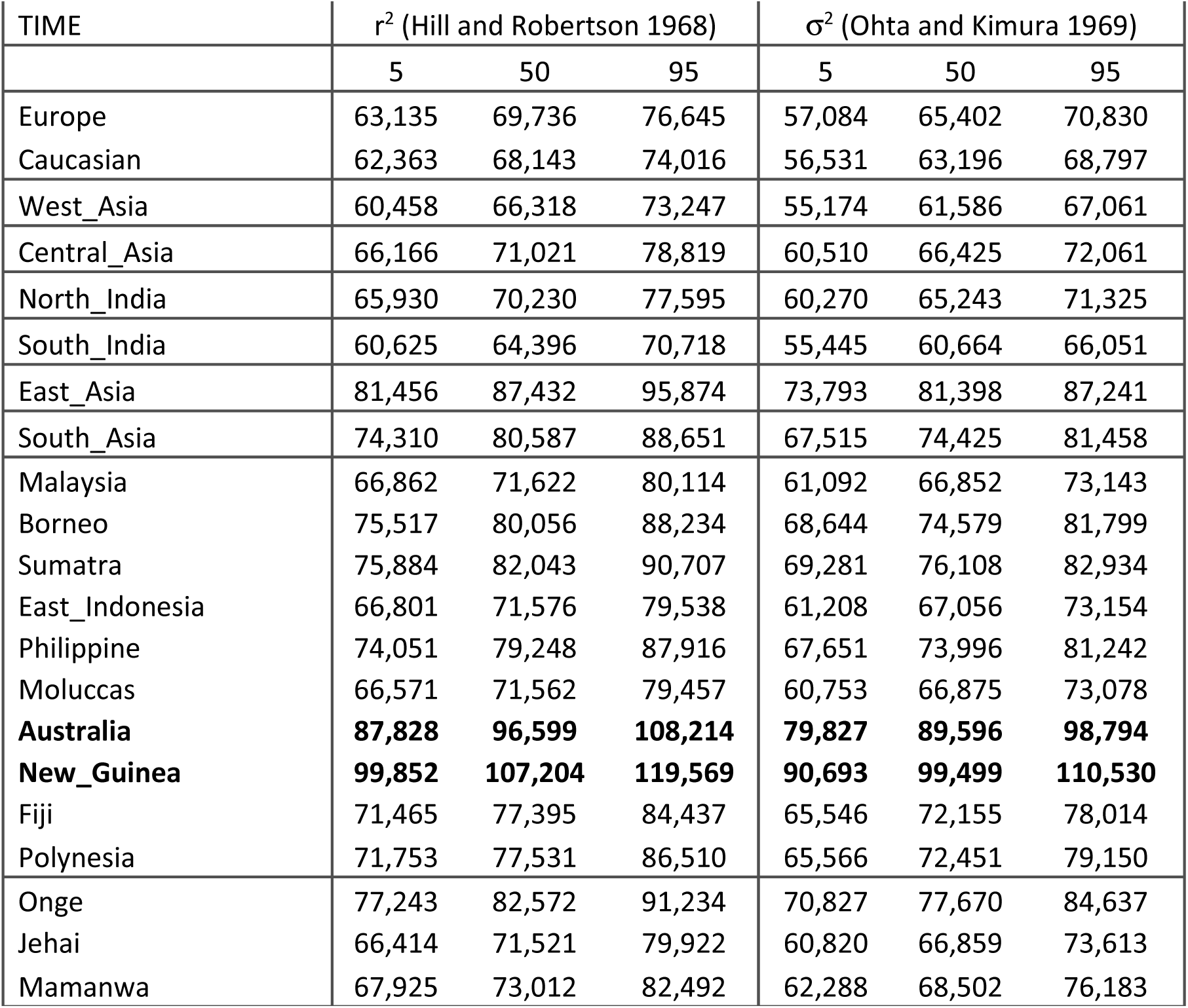
Estimated divergence times from (East) Africa using the r^2^ or σ^2^ statistics as estimator of LD level. For each comparisons with East African, the three columns report the 95% lower confidence limit, the point estimate (in years, assuming a generation interval =25 years [85]), and the 95% upper confidence limit.

| TIME | $r^2$ (Hill and Robertson 1968) | | | $\sigma^2$ (Ohta and Kimura 1969) | | |
| --- | --- | --- | --- | --- | --- | --- |
|  | 5 | 50 | 95 | 5 | 50 | 95 |
| Europe | 63,135 | 69,736 | 76,645 | 57,084 | 65,402 | 70,830 |
| Caucasian | 62,363 | 68,143 | 74,016 | 56,531 | 63,196 | 68,797 |
| West_Asia | 60,458 | 66,318 | 73,247 | 55,174 | 61,586 | 67,061 |
| Central_Asia | 66,166 | 71,021 | 78,819 | 60,510 | 66,425 | 72,061 |
| North_India | 65,930 | 70,230 | 77,595 | 60,270 | 65,243 | 71,325 |
| South_India | 60,625 | 64,396 | 70,718 | 55,445 | 60,664 | 66,051 |
| East_Asia | 81,456 | 87,432 | 95,874 | 73,793 | 81,398 | 87,241 |
| South_Asia | 74,310 | 80,587 | 88,651 | 67,515 | 74,425 | 81,458 |
| Malaysia | 66,862 | 71,622 | 80,114 | 61,092 | 66,852 | 73,143 |
| Borneo | 75,517 | 80,056 | 88,234 | 68,644 | 74,579 | 81,799 |
| Sumatra | 75,884 | 82,043 | 90,707 | 69,281 | 76,108 | 82,934 |
| East_Indonesia | 66,801 | 71,576 | 79,538 | 61,208 | 67,056 | 73,154 |
| Philippine | 74,051 | 79,248 | 87,916 | 67,651 | 73,996 | 81,242 |
| Moluccas | 66,571 | 71,562 | 79,457 | 60,753 | 66,875 | 73,078 |
| **Australia** | **87,828** | **96,599** | **108,214** | **79,827** | **89,596** | **98,794** |
| **New_Guinea** | **99,852** | **107,204** | **119,569** | **90,693** | **99,499** | **110,530** |
| Fiji | 71,465 | 77,395 | 84,437 | 65,546 | 72,155 | 78,014 |
| Polynesia | 71,753 | 77,531 | 86,510 | 65,566 | 72,451 | 79,150 |
| Onge | 77,243 | 82,572 | 91,234 | 70,827 | 77,670 | 84,637 |
| Jehai | 66,414 | 71,521 | 79,922 | 60,820 | 66,859 | 73,613 |
| Mamanwa | 67,925 | 73,012 | 82,492 | 62,288 | 68,502 | 76,183 |

### Comparing the predictions of single vs multiple African exit models: Divergence times

Having shown that significantly different times of separation from Africa are estimated for Europe and Australia/New Guinea, the question arises whether it would be possible to obtain such results by chance alone, had AMH dispersed in a single wave, at the time period at which that dispersal is generally placed (in the calculations that follow, we always considered the *N*_e_ and *T* estimates obtained using the unweighted *r*^2^ statistic). To answer, we needed a null distribution of *T* values under the SD model, which we constructed by simulation, using the software *ms* [56]. We plotted the (null) distribution of the 24,000 separation times derived from the simulations and we compared it with the observed *T* estimates. Whereas the value estimated in the European sample falls perfectly within the range of times predicted by the classical SD model, that is not the case for the New Guinean and the Australian values, falling in the right tail of this distribution at *P*<0.05 level (Fig. 4). This can only mean that a single exit from Africa, even considering the uncertainty in our knowledge of the relevant parameters, cannot account for the differences in the separation times from Africa observed, respectively, in Europe on the one hand, and in Australo-Melanesia on the other.

**Figure 4.**
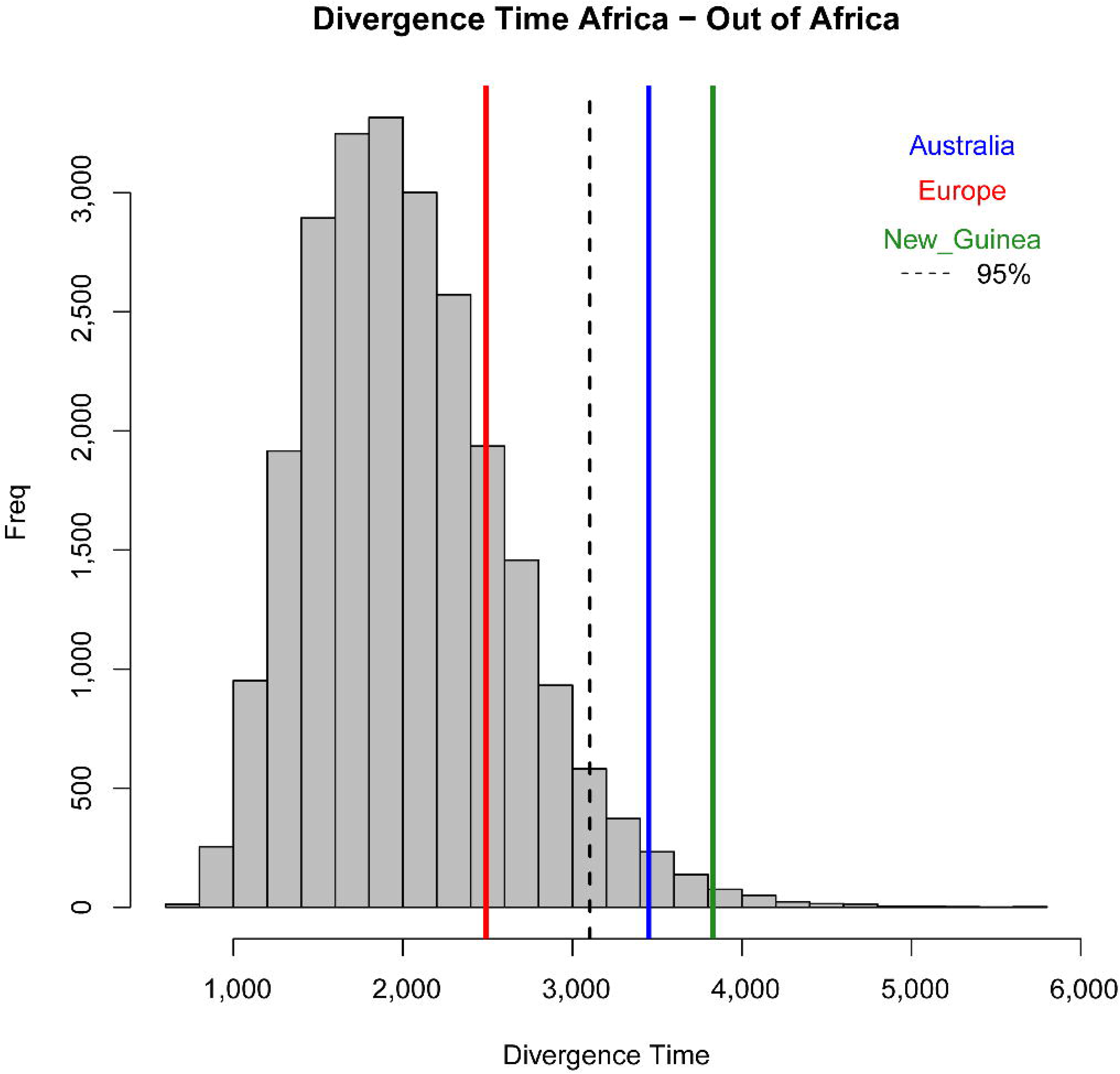
Comparison of three observed divergence times with the distribution of 24,000 divergence times between African and non-African populations generated by simulation of a SD model. Data generated for 24 combinations of effective population sizes (3,000 ≤ *N*_e_ ≤ 8,000) and divergence times (40 k years ago ≤ *T* ≤ 70 k years ago), 1,000 independent datasets for each such combination. At every iteration, genetic variation at 1Mb was considered in 100 chromosomes per population, thus analyzing 200,000 Mb for each parameter combination (for a total of 4,800 Gb in 24,000 iterations, see Supplemental Methods for details).

### Possible effects of a Denisovan admixture in Melanesia

Recent analyses of the genetic relationships between modern humans and Denisovans suggested that a fraction possibly as high as 6-8% of the Melanesian genomes may be traced back to Denisovan ancestor [60]. To rule out the possibility that the apparent difference in African divergence times for Europe and Australo-Melanesia may somewhat reflect Denisovan admixture, we removed from the analysis the SNPs that were identified as representing the Denisovan contribution to the latter’s genome. We recalculated the *F*_ST_s from the 80,621 SNPs obtained from the filtering process, and reestimated the divergence times from Africa, finding they are still very close to those previously estimated (Additional File 10 and 11).

### Estimates of population admixture

Other Far Eastern populations, besides Australia and New Guineans for which we estimated a remote separation from Africans, may have taken part in an early exit from Africa through a Southern route. Identifying them is not straightforward, though, because we basically have a continuous set of divergence times from East Africa, from 66K to 107K years ago (Table 1). This result is consistent with both a continuous migration process from Africa across some 40K years (which so far has never been proposed, to the best of our knowledge) and with an early exit, followed by genetic exchanges with later-dispersing groups, which has diluted or erased altogether the genetic evidence of the earliest migration. Our previous ADMIXTURE analysis highlighted an ancestral genetic component (green, Figure 3) to which all Australo-Melanesian genotypes could be associated. In what follows, we explored the possibility that the same component be a marker of the earliest African exit in other populations as well. To understand whether that could have actually been the case, we used a method, *TreeMix* [62] to estimate from genome-wide data a maximum-likelihood tree of populations, and then to infer events of gene flow after the split by identifying populations that poorly fit the tree; if admixture was extensive we expect to observe extensive reticulation in the tree. We selected from our dataset just the populations showing at least 30% of the green Admixture component at *k*=5 and clustering together in the third group of the DAPC scatter plot, choosing the East Asia sample as outgroup. Additional File 12 shows the maximum-likelihood tree. Evidence for extensive genetic exchanges after population splits is apparent from East Asia (represented in light blue in the tree) toward populations putatively involved in the early African dispersal (represented in green in the tree). Prior to adding these migration episodes the graph captures 87% of the global variance in the data; including the top 6 migration events (indicated by arrows coloured according to the intensities of the process) brought this percentage to 99% (Additional File 13). Therefore, these results support the hypothesis that relatively recent admixture events could have obscured the genomic signatures of the first migration out of Africa in these Southeast Asian populations, ultimately biasing downwards the estimates of their divergence times from Africans.

### Comparing the predictions of single vs multiple African exit models: Geographical patterns

To conclude, we tried to better define some details of the AMH dispersal out of the African continent by evaluating which geographical migration route can better account for the current patterns of genome diversity. To minimize the effects of recent gene flow unrelated with the first human dispersals, which was clearly not negligible (see previous section) we selected populations with at least 80% of a single ancestral component in the ADMIXTURE results (i.e. Australia, the Caucasus, East Africa, East Asia, Europe, New Guinea, South Africa, South India, West Africa). All the Mantel correlations thus calculated were positive and significant (Table 2), suggesting that all tested models succeed in plausibly predicting the observed patterns of genome diversity. The highest correlation observed for Model 3 (*r*=0.767) supports the southern route hypothesis for populations of South-East Asia and Oceania, but the difference between Models 3 and 1 is not significant by Fisher’s criterion [68] (*Z*=-1.26, *P*=0.08).

**Table 2.** Partial Mantel correlations between genetic and geographic distances. Comparisons of the genetic distance matrix (*F*_ST_) with the geographic distances calculated according to the three dispersal models, while holding constant population divergence values (*T*). Values are Pearson correlation coefficients, and the *P*-values have been empirically calculated over 10,000 permutations of one matrix’ rows and columns.

|  | Partial Mantel Test |  |
| --- | --- | --- |
|  | r | p-value |
| **Model 1** | 0.67 | 0.0001 |
| **Model 2** | 0.64 | 0.0012 |
| **Model 3** | **0.77** | 0.0001 |

### Discussion

Establishing whether genomic differences among populations are compatible with a single, major expansion from Africa is crucial for reconstructing the set of migration processes leading to the human peopling of the Old World. Two factors, namely the effects of population sizes and of admixture after the main population split, complicate the estimation of divergence times between populations. Past population sizes are unknown, and are generally estimated from genetic diversity, under neutrality assumptions. However, genetic differences between populations are large if the populations long evolved independently, or if they had small effective sizes, or by any combination of these factors. To circumvent this problem, in this study we resorted to *LD* values to estimate long-term population sizes and separate their effect from that of population history. This way, we found that the populations at the extremes of the geographical range considered differ substantially in the timing of their separation from the Eastern African populations. This difference is statistically significant, and we showed by simulation that it cannot possibly be reconciled with a model assuming a single, major dispersal of all non-Africans through the classical Northern route (as pointed by Pagani et al.[69]). The model we tested is necessarily simple and does not take into account potential admixture with archaic human forms. However, since the estimated degree of Neandertal ancestry is the same in all modern non-African populations [4], the inclusion of this event would only affect the *absolute* values of divergence times from Africa, and not the *ratio* between them. Conversely, Denisovan admixture may inflate the estimated divergence times in the Easternmost populations. Removal of the SNPs identified as a potential Denisovan contribution to the modern genomes caused no substantial change in the results, showing that our estimates reflect to a minimal extent, if any, the effects of interbreeding with Denisovans [60].

As for admixture among modern populations after the split from Africa, which may affect estimates of their divergence time [70], the demographic history of large sections of Asia is too complex and elusive to allow one to tell apart admixed and non-admixed groups. However, we argue the impact of modern admixture upon our results cannot be too strong, because geographically-intermediate populations were excluded from the final analysis. This way, significant differences in time estimates were observed for populations (Europe, the Caucasus, New Guinea and Australia) showing a rather homogeneous genetic composition in the ADMIXTURE analysis, with most individual genotypes attributed to a single ancestral component (Figure 3).

The method used in the present study allows us to safely rule out that fluctuations in long-term population sizes might have distorted our time estimates. Three-fold differences in very ancient (e.g. > 100,000 years ago; Additional File 7) population sizes may appear, at a first sight, difficult to justify, because at that time all *N*_e_ values should converge to a value representing the size of the common ancestral African population. However, a similar result was also obtained in the only previous study based on the same method [18], and interpreted as reflecting founder effects accompanying the dispersal from Africa. In turn, these phases of increased genetic drift may have increased *LD*, and hence caused underestimation of *N*_e_ in all non-Africans. However, the resulting distortion, if any, should have affected the *absolute* values of *T*, but not the *relative* timing of the Europeans’ and Asians’ separation from Africans, which is what this study is concerned with. Another possibility is that 100,000 or so years ago the ancestors of current Eurasians were already genetically distinct from the ancestors of modern Africans (as proposed by Refs. [71, 72]). If so, the different *N*_e_ estimates of the present study would not be a statistical artifact, but would reflect actual differences between geographically-isolated ancient populations.

Two independent analyses (by ADMIXTURE and *TreeMix*) suggest that the genotypes of most Central Asians reflect variable degrees of gene flow between populations which may have left Africa in different waves. As a result, the distribution of divergence times is essentially continuous, and hence it would make no sense to try to classify Central Asian populations as derived from either the first or the second African exit under the model of multiple dispersals.

When we modeled population dispersal in space, the correlation between genetic and geographic distances was higher under the MD than under the SD model, but this difference was statistically insignificant (Table 2). This seems likely due to the fact that the three models being compared share several features, such as the same set of geographic/genetic distances for the European populations, which reduces the power of any test. However, the separation times previously estimated made us confident that the SD model is inconsistent with the data, and so what was really important at this stage was the comparison between the two MD models. The better fit of model 3 than model 2 implies that the MD model works better if only part of the Asian genomic diversity is attributed to the earliest dispersal. A better fit of a MD than of a SD model was also observed in parallel analyses of cranial measures and of a much smaller genomic dataset [13], suggesting that our findings may indeed reflect a general pattern of human diversity.

The data we analyzed are probably affected, to an unknown but not negligible extent, by a bias due to the fact that most SNPs in the genotyping platforms were discovered in European populations; however the measure we used to calculate *N*_e_ and hence the separation time, *LD*, is expected to be relatively unaffected by this kind of bias [18, 73]. At any rate, a likely effect of such a bias would be a spurious increase of the estimated differences between Europeans and the populations being compared with them, Africans in this case. Quite to the contrary, here the Europeans appeared significantly *closer* to Africans than Australo-Melanesians, a result which therefore cannot be due to that kind of ascertainment bias.

Can selection account, at least in part, for these findings? In principle, we have no way to rule this out. However, in practice, even though positive selection may have extensively affected the human genome, large allele-frequency shifts at individual loci are surprisingly rare [74], so much so that so far only for very few SNPs any effects of selection have been demonstrated[75]. If we also consider that genomic regions with large allele-frequency differences are not generally associated with high levels of linkage disequilibrium, in contrast with what would be expected after a selective sweep [74, 76], it seems fair to conclude that the main allele frequency shifts occurred in a rather remote past and are unlikely to reflect geographic differences in the selection regimes [77]. In any case, only 8% of the SNPs we considered map within expressed loci, or in their control regions (Figure 5); therefore, the impact of selection upon the results of this study, if any, can hardly be regarded as substantial.

**Figure 5.**
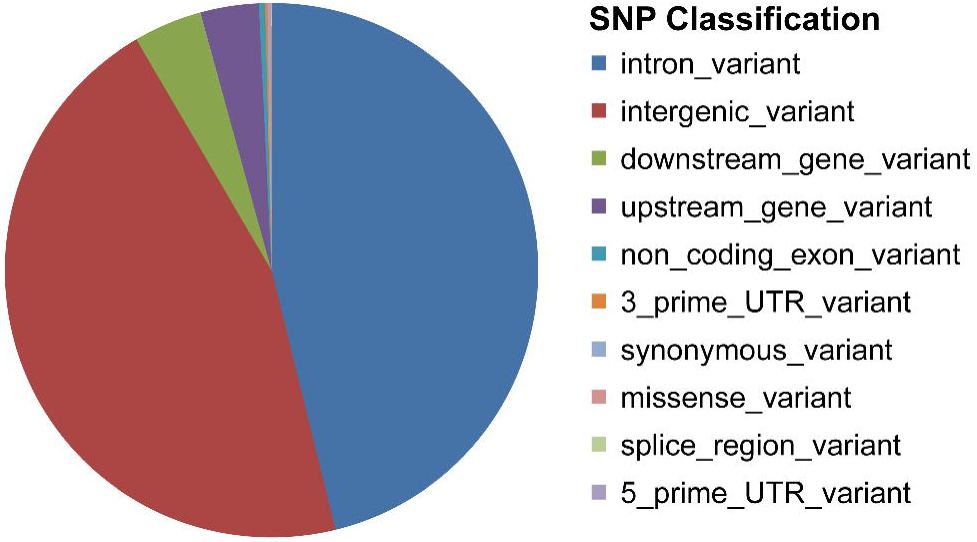
Distribution of the SNPs considered in functionally different genome regions.

## Conclusions

Analyses of genomic data based on different sets of assumptions and different methods agree in indicating: (i) that a model with a single early dispersal from Africa fails to account for one crucial aspect of human genome diversity, the distribution of divergence times from Africa, and (ii) that within the model of multiple dispersal, geographical patterns of genome diversity are more accurately predicted assuming that not all Asian and New Guinea/Australian populations have had the same evolutionary history.

In the light of these results, we propose that at least two major dispersal phenomena from Africa led to the peopling of Eurasia and Australo-Melanesia. These phenomena seem clearly distinct both in their timing and in their geographical scope.

The view whereby only part of the ancestors of current non-African populations dispersed through the Levant has some non-trivial consequences upon the possible interactions between AMH and archaic forms, traces of whose genomes have been identified in many non-African populations, including New Guineans [4, 78]. The estimated contribution of Neandertals is less in the European than in the Asian/Melanesian genomes, despite the long coexistence between Neandertals and Europeans [79]. At present, the standard way to explain this finding is to assume one single, major episode of hybridization in Palestine (or perhaps further North and East [80]) 47K to 65K years ago [58], followed by a split between the Europeans’ ancestors on the one hand, and the Asians’ and Oceanians’ on the other [80, 81]. After that, additional contacts might have occurred, but only between Neandertals and Asians [82]. However, if most ancestors of New Guineans dispersed through a Southern route, as this study shows, they would have missed by 2,000 km or so the nearest documented Neandertals with whom they could have intercrossed. Thus, this study raises the possibility that the current patterns of human diversity need more complex models to be fully explained. One possibility is that admixture with Neandertals might have occurred before AMH left Africa [83]. Another is that common ancestry, rather than hybridization, may account for the excess similarity of Eurasians with Neandertals, in the presence of an ancient structuring of populations [72, 84]. A third possibility is that the apparent traces of Neandertal hybridization in Papua New Guinea may in fact be due to Denisovan admixture. We are not in a position to actually test for these possibilities, but exploring these hypotheses may contribute to a better understanding of the main human dispersal processes and of the relationships between archaic and modern human forms.

## Competing financial interests

All authors declare no competing financial interests.

## Author contributions

S.G., F.T. and G.B. conceived and designed the analyses; S.G., F.T., M.M.,S.V.T. and L.D.S. analyzed the data; G.B., S.G. and F.T wrote the manuscript.

## Acknowledgements

This study was supported by the European Research Council ERC-2011-AdG_295733 grant (LanGeLin), and in part by a grant of the Italian Ministry for Research and Universities (MIUR) PRIN 2010-2011. We are indebted to Maria Teresa Vizzari for technical help, and to Andrea Benazzo, Cesare de Filippo and Johannes Krause for suggestions and critical discussion of this manuscript.

## Additional Files

### Additional File 1

Format: .tif

Geographic location of all the 71 populations analyzed, the different dataset we use are represented by different colors and are detailed in Additional File 2.

### Additional File 2

Format: .pdf

General information about the 24 metapopulations analyzed.

### Additional File 3

Format: .pdf

Representation of the human demographic model tested by ms. The past is at the top, the present is at the bottom.

### Additional File 4

Format: .pdf

Estimation of the most likely number of clusters in the data (X-axis) as a function of the cross-validation error observed in the attempted assignments (Y-axis).

### Additional File 5

Format: .pdf

Inference of the most likely number of clusters in the DAPC. A *k* value of 6 (the lowest BIC value) represents the best summary of the data.

### Additional File 6

Format: .pdf

Discriminant Analysis of Principal Components, DAPC. (a) classification of individual genotypes; for each row (each population) the figures refer to the numbers of individuals assigned to of the k=6 clusters, each cluster associated with a different colour; (b) scatterplot along the first two axes; each symbol corresponds to an individual genotypes; in the insets, the fraction of Principal Components retained in the analysis (left) and the fraction of the overall variance attributed to the first five eigenvalues, with the first two columns, in grey, representing the first two Discriminant Functions (right).

### Additional File 7

Format: .pdf

Estimates of Ne from measures of linkage disequilibrium, using the (a) r2 and (b) σ2 statistics as estimator of LD level. Time is on the X-axis and is expressed in generations from the present. Very recent estimates have been omitted because not reliably estimated.

### Additional File 8

Format: .pdf

Harmonic means of the estimated population effective sizes (*N*_*e*_), using the (a) r^2^ and (b) σ^2^ statistics as estimator of LD level. Vertical bars represent empirical 90% confidence estimates.

### Additional File 9

Format: .pdf

Pairwise FST values estimated between populations. The matrix is symmetrical.

### Additional File 10

Format: .pdf

Estimates of population divergence times, using (a) r^2^ and (b) σ^2^ statistics, as estimator of LD level.

### Additional File 11

Format: .pdf

Population divergence time estimated on a subset of SNPs chosen to exclude the effect of an archaic introgression from Denisovan.

### Additional File 12

Format: .pdf

Population relationships inferred by *TreeMix*. The Maximum-likelihood tree is in black; branch lengths are proportional to the impact of genetic drift, which may or may not faithfully represent separation times between populations. The inferred migration events are represented by arrows pointing from the putative source to the putative target populations, with colours of the arrows representing the relative weight of the genetic exchanges, according to the heat scale on the left.

### Additional File 13

Format: .pdf

Fractions of the total variance explained by the model at increasing numbers of migrations superimposed to the bifurcating tree in the *TreeMix* analysis.\

## References

1. Aubert M, Pike AW, Stringer C, Bartsiokas A, Kinsley L, Eggins S et al. Confirmation of a late middle Pleistocene age for the Omo Kibish 1 cranium by direct uranium-series dating. J Hum Evol. 2012;63(5):704–10.

2. McDougall I, Brown FH, Fleagle JG. Stratigraphic placement and age of modern humans from Kibish, Ethiopia. Nature. 2005;433(7027):733–6.

3. Stringer C. Modern human origins: progress and prospects. Philos Trans R Soc Lond B Biol Sci. 2002;357(1420):563–79.

4. Green RE, Krause J, Briggs AW, Maricic T, Stenzel U, Kircher M et al. A draft sequence of the Neandertal genome. Science. 2010;328(5979):710–22.

5. Reich D, Patterson N, Kircher M, Delfin F, Nandineni MR, Pugach I et al. Denisova admixture and the first modern human dispersals into Southeast Asia and Oceania. Am J Hum Genet. 2011;89(4):516–28.

6. Hammer MF, Woerner AE, Mendez FL, Watkins JC, Wall JD. Genetic evidence for archaic admixture in Africa. Proc Natl Acad Sci U S A. 2011;108(37):15123–8.

7. Lachance J, Vernot B, Elbers CC, Ferwerda B, Froment A, Bodo JM et al. Evolutionary history and adaptation from high-coverage whole-genome sequences of diverse African hunter-gatherers. Cell. 2012;150(3):457–69.

8. Abi-Rached L, Jobin MJ, Kulkarni S, McWhinnie A, Dalva K, Gragert L et al. The shaping of modern human immune systems by multiregional admixture with archaic humans. Science. 2011;334(6052):89–94.

9. Khrameeva EE, Bozek K, He L, Yan Z, Jiang X, Wei Y et al. Neanderthal ancestry drives evolution of lipid catabolism in contemporary Europeans. Nat Commun. 2014;5:3584.

10. Liu N, Zhao H. A non-parametric approach to population structure inference using multilocus genotypes. Hum Genomics. 2006;2(6):353–64.

11. Fu Q, Mittnik A, Johnson PL, Bos K, Lari M, Bollongino R et al. A revised timescale for human evolution based on ancient mitochondrial genomes. Curr Biol. 2013;23(7):553–9.

12. Lahr MM, Foley RA. Multiple Dispersals and Modern Human Origins. Evolutionary Anthropology. 1994;3:48–60.

13. Reyes-Centeno H, Ghirotto S, Detroit F, Grimaud-Herve D, Barbujani G, Harvati K. Genomic and cranial phenotype data support multiple modern human dispersals from Africa and a southern route into Asia. Proc Natl Acad Sci U S A. 2014;111(20):7248–53.

14. Field JS, Petraglia MD, Lahr MM. The southern dispersal hypothesis and the South Asian archaeological record: Examination of dispersal routes through GIS analysis. Journal of Anthropological Archaeology. 2007;26:88–108.

15. Di D, Sanchez-Mazas A. Challenging views on the peopling history of East Asia: the story according to HLA markers. Am J Phys Anthropol. 2011;145(1):81–96.

16. Ghirotto S, Penso-Dolfin L, Barbujani G. Genomic evidence for an African expansion of anatomically modern humans by a Southern route. Hum Biol. 2011;83(4):477–89.

17. Macaulay V, Hill C, Achilli A, Rengo C, Clarke D, Meehan W et al. Single, rapid coastal settlement of Asia revealed by analysis of complete mitochondrial genomes. Science. 2005;308(5724):1034–6.

18. McEvoy BP, Powell JE, Goddard ME, Visscher PM. Human population dispersal “Out of Africa” estimated from linkage disequilibrium and allele frequencies of SNPs. Genome Res. 2011;21(6):821–9.

19. Quintana-Murci L, Semino O, Bandelt HJ, Passarino G, McElreavey K, Santachiara-Benerecetti AS. Genetic evidence of an early exit of Homo sapiens sapiens from Africa through eastern Africa. Nat Genet. 1999;23(4):437–41.

20. Rasmussen M, Guo X, Wang Y, Lohmueller KE, Rasmussen S, Albrechtsen A et al. An Aboriginal Australian genome reveals separate human dispersals into Asia. Science. 2011;334(6052):94–8.

21. Prugnolle F, Manica A, Balloux F. Geography predicts neutral genetic diversity of human populations. Curr Biol. 2005;15(5):R159–60.

22. Li JZ, Absher DM, Tang H, Southwick AM, Casto AM, Ramachandran S et al. Worldwide human relationships inferred from genome-wide patterns of variation. Science. 2008;319(5866):1100–4.

23. DeGiorgio M, Jakobsson M, Rosenberg NA. Out of Africa: modern human origins special feature: explaining worldwide patterns of human genetic variation using a coalescent-based serial founder model of migration outward from Africa. Proc Natl Acad Sci U S A. 2009;106(38):16057–62.

24. Tishkoff SA, Goldman A, Calafell F, Speed WC, Deinard AS, Bonne-Tamir B et al. A global haplotype analysis of the myotonic dystrophy locus: implications for the evolution of modern humans and for the origin of myotonic dystrophy mutations. Am J Hum Genet. 1998;62(6):1389–402.

25. Deshpande O, Batzoglou S, Feldman MW, Cavalli-Sforza LL. A serial founder effect model for human settlement out of Africa. Proc Biol Sci. 2009;276(1655):291–300.

26. Ramachandran S, Deshpande O, Roseman CC, Rosenberg NA, Feldman MW, Cavalli-Sforza LL. Support from the relationship of genetic and geographic distance in human populations for a serial founder effect originating in Africa. Proc Natl Acad Sci U S A. 2005;102(44):15942–7.

27. Gonzalez-Ruiz M, Santos C, Jordana X, Simon M, Lalueza-Fox C, Gigli E et al. Tracing the origin of the east-west population admixture in the Altai region (Central Asia). PLoS One. 2012;7(11):e48904.

28. Comas D, Plaza S, Wells RS, Yuldaseva N, Lao O, Calafell F et al. Admixture, migrations, and dispersals in Central Asia: evidence from maternal DNA lineages. Eur J Hum Genet. 2004;12(6):495–504.

29. Martinez-Cruz B, Vitalis R, Segurel L, Austerlitz F, Georges M, Thery S et al. In the heartland of Eurasia: the multilocus genetic landscape of Central Asian populations. Eur J Hum Genet. 2011;19(2):216–23.

30. Nievergelt CM, Maihofer AX, Shekhtman T, Libiger O, Wang X, Kidd KK et al. Inference of human continental origin and admixture proportions using a highly discriminative ancestry informative 41-SNP panel. Investig Genet. 2013;4(1):13.

31. Mezzavilla M, Vozzi D, Pirastu N, Girotto G, d’Adamo P, Gasparini P et al. Genetic landscape of populations along the Silk Road: admixture and migration patterns. BMC Genet. 2014;15(1):131.

32. Lopez Herraez D, Bauchet M, Tang K, Theunert C, Pugach I, Li J et al. Genetic variation and recent positive selection in worldwide human populations: evidence from nearly 1 million SNPs. PLoS One. 2009;4(11):e7888.

33. Pugach I, Delfin F, Gunnarsdottir E, Kayser M, Stoneking M. Genome-wide data substantiate Holocene gene flow from India to Australia. Proc Natl Acad Sci U S A. 2013;110(5):1803–8.

34. Reich D, Thangaraj K, Patterson N, Price AL, Singh L. Reconstructing Indian population history. Nature. 2009;461(7263):489–94.

35. Xing J, Watkins WS, Witherspoon DJ, Zhang Y, Guthery SL, Thara R et al. Fine-scaled human genetic structure revealed by SNP microarrays. Genome Res. 2009;19(5):815–25.

36. Xing J, Watkins WS, Shlien A, Walker E, Huff CD, Witherspoon DJ et al. Toward a more uniform sampling of human genetic diversity: a survey of worldwide populations by high-density genotyping. Genomics. 2010;96(4):199–210.

37. Purcell S, Neale B, Todd-Brown K, Thomas L, Ferreira MA, Bender D et al. PLINK: a tool set for whole-genome association and population-based linkage analyses. Am J Hum Genet. 2007;81(3):559–75.

38. Relethford JH. Examination of the relationship between inbreeding and population size. J Biosoc Sci. 1985;17(1):97–106.

39. R Development Core Team. R: A Language and Environment for Statistical Computing. Vienna, Austria: the R Foundation for Statistical Computing. ISBN: 3-900051-07-0 Available online at http://wwwR-projectorg/. 2011

40. Alexander DH, Novembre J, Lange K. Fast model-based estimation of ancestry in unrelated individuals. Genome Res. 2009;19(9):1655–64.

41. Jakobsson M, Rosenberg NA. CLUMPP: a cluster matching and permutation program for dealing with label switching and multimodality in analysis of population structure. Bioinformatics. 2007;23(14):1801–6.

42. Rosenberg NA. Distruct: a program for the graphical display of population Molecular Ecology Notes 2004;4:137–8.

43. Jombart T, Devillard S, Balloux F. Discriminant analysis of principal components: a new method for the analysis of genetically structured populations. BMC Genet. 2010;11:94.

44. Jombart T, Ahmed I. adegenet 1.3-1: new tools for the analysis of genome-wide SNP data. Bioinformatics. 2011;27(21):3070–1.

45. Weir BS, Cockerham CC. Estimating F-statistics for the analysis of population structure. Evolution 1984;38:1358–70.

46. Tenesa A, Navarro P, Hayes BJ, Duffy DL, Clarke GM, Goddard ME et al. Recent human effective population size estimated from linkage disequilibrium. Genome Res. 2007;17(4):520–6.

47. Hayes BJ, Visscher PM, McPartlan HC, Goddard ME. Novel multilocus measure of linkage disequilibrium to estimate past effective population size. Genome Res. 2003;13(4):635–43.

48. Clark AG, Hubisz MJ, Bustamante CD, Williamson SH, Nielsen R. Ascertainment bias in studies of human genome-wide polymorphism. Genome Res. 2005;15(11):1496–502.

49. Hill WG, Robertson A. Linkage disequilibrium in finite populations. Theor Appl Genet. 1968;38(6):226–31.

50. Ohta T, Kimura M. Linkage disequilibrium at steady state determined by random genetic drift and recurrent mutation. Genetics. 1969;63(1):229–38.

51. Sved JA. Linkage disequilibrium and homozygosity of chromosome segments in finite populations. Theor Popul Biol. 1971;2(2):125–41.

52. McVean GA. A genealogical interpretation of linkage disequilibrium. Genetics. 2002;162(2):987–91.

53. Holsinger KE, Weir BS. Genetics in geographically structured populations: defining, estimating and interpreting F(ST). Nat Rev Genet. 2009;10(9):639–50.

54. Mezzavilla M, Ghirotto S. Neon: An R Package to Estimate Human Effective Population Size and Divergence Time from Patterns of Linkage Disequilibrium between SNPS. J Comput Sci Syst Biol 2015(8 4.):037–04.

55. Benazzo A, Panziera A, Bertorelle G. 4P: fast computing of population genetics statistics from large DNA polymorphism panels. Ecology and Evolution. 2014;(in press)

56. Hudson RR. Generating samples under a Wright-Fisher neutral model of genetic variation. Bioinformatics. 2002;18(2):337–8.

57. Mellars P, Gori KC, Carr M, Soares PA, Richards MB. Genetic and archaeological perspectives on the initial modern human colonization of southern Asia. Proc Natl Acad Sci U S A. 2013;110(26):10699–704.

58. Sankararaman S, Patterson N, Li H, Paabo S, Reich D. The date of interbreeding between Neandertals and modern humans. PLoS Genet. 2012;8(10):e1002947.

59. Petraglia MD, Haslam M, Fuller DQ, Boivin N, Clarkson C. Out of Africa: new hypotheses and evidence for the dispersal of Homo sapiens along the Indian Ocean rim. Annals of human biology. 2010;37(3):288–311.

60. Meyer M, Kircher M, Gansauge MT, Li H, Racimo F, Mallick S et al. A high-coverage genome sequence from an archaic Denisovan individual. Science. 2012;338(6104):222–6.

61. Danecek P, Auton A, Abecasis G, Albers CA, Banks E, DePristo MA et al. The variant call format and VCFtools. Bioinformatics. 2011;27(15):2156–8.

62. Pickrell JK, Pritchard JK. Inference of population splits and mixtures from genome-wide allele frequency data. PLoS Genet. 2012;8(11):e1002967.

63. McRae BH. Isolation by resistance. Evolution. 2006;60(8):1551–61.

64. Oppenheimer S. A single southern exit of modern humans from Africa: Before or after Toba?. Quaternary International 2012;258:88–9.

65. Beyin A. Upper Pleistocene Human Dispersals out of Africa: A Review of the Current State of the Debate. Int J Evol Biol. 2011;2011:615094.

66. Mantel N. The detection of disease clustering and a generalized regression approach. Cancer Res. 1967;27(2):209–20.

67. Henn BM, Botigue LR, Gravel S, Wang W, Brisbin A, Byrnes JK et al. Genomic ancestry of North Africans supports back-to-Africa migrations. PLoS Genet. 2012;8(1):e1002397.

68. Fisher RA. On the probable error of a coefficient of correlation deduced from a small sample. Metron. 1921;1:3–32.

69. Pagani L, Schiffels S, Gurdasani D, Danecek P, Scally A, Chen Y et al. Tracing the Route of Modern Humans out of Africa by Using 225 Human Genome Sequences from Ethiopians and Egyptians. Am J Hum Genet. 2015;96(6):986–91.

70. Sved JA. Correlation measures for linkage disequilibrium within and between populations. Genet Res (Camb). 2009;91(3):183–92.

71. Harding RM, McVean G. A structured ancestral population for the evolution of modern humans. Curr Opin Genet Dev. 2004;14(6):667–74.

72. Eriksson A, Manica A. Effect of ancient population structure on the degree of polymorphism shared between modern human populations and ancient hominins. Proc Natl Acad Sci U S A. 2012;109(35):13956–60.

73. Jakobsson M, Scholz SW, Scheet P, Gibbs JR, VanLiere JM, Fung HC et al. Genotype, haplotype and copy-number variation in worldwide human populations. Nature. 2008;451(7181):998–1003.

74. Coop G, Pickrell JK, Novembre J, Kudaravalli S, Li J, Absher D et al. The role of geography in human adaptation. PLoS Genet. 2009;5(6):e1000500.

75. Hernandez RD, Kelley JL, Elyashiv E, Melton SC, Auton A, McVean G et al. Classic selective sweeps were rare in recent human evolution. Science. 2011;331(6019):920–4.

76. Pritchard JK, Pickrell JK, Coop G. The genetics of human adaptation: hard sweeps, soft sweeps, and polygenic adaptation. Curr Biol. 2010;20(4):R208–15.

77. Alves I, Sramkova Hanulova A, Foll M, Excoffier L. Genomic data reveal a complex making of humans. PLoS Genet. 2012;8(7):e1002837.

78. Sankararaman S, Mallick S, Dannemann M, Prufer K, Kelso J, Paabo S et al. The genomic landscape of Neanderthal ancestry in present-day humans. Nature. 2014;507(7492):354–7.

79. Higham T, Douka K, Wood R, Ramsey CB, Brock F, Basell L et al. The timing and spatiotemporal patterning of Neanderthal disappearance. Nature. 2014;512(7514):306–9.

80. Prufer K, Racimo F, Patterson N, Jay F, Sankararaman S, Sawyer S et al. The complete genome sequence of a Neanderthal from the Altai Mountains. Nature. 2014;505(7481):43–9.

81. Stoneking M, Krause J. Learning about human population history from ancient and modern genomes. Nat Rev Genet. 2011;12(9):603–14.

82. Vernot B, Akey JM. Resurrecting surviving Neandertal lineages from modern human genomes. Science. 2014;343(6174):1017–21.

83. Sanchez-Quinto F, Botigue LR, Civit S, Arenas C, Avila-Arcos MC, Bustamante CD et al. North African populations carry the signature of admixture with Neandertals. PLoS One. 2012;7(10):e47765.

84. Ray N, Currat M, Berthier P, Excoffier L. Recovering the geographic origin of early modern humans by realistic and spatially explicit simulations. Genome Res. 2005;15(8):1161–7.

85. Fenner JN. Cross-cultural estimation of the human generation interval for use in genetics-based population divergence studies. Am J Phys Anthropol. 2005;128(2):415–23.

